# The host protein cyclophilin A inhibits HIV-1 nuclear entry by decreasing capsid elasticity

**DOI:** 10.1101/2025.09.03.673798

**Authors:** Jun Hong, Akshay Deshpande, Yatish Thakare, Lora Simonovsky, AidanDarian W. Douglas, Conall Mc Guinness, Noa Rotem-Dai, Michelle L. Kortyna, J Ole Klarhof, Jiong Shi, Till Boecking, Leo C James, Ashwanth C Francis, Itay Rousso, Christopher Aiken

## Abstract

Binding of the host protein cyclophilin A (CypA) to the HIV-1 capsid exerts a variety of effects on infection, including enhancement of reverse transcription, stabilization of the capsid, and promotion of nuclear entry. For several HIV-1 mutants, CypA binding inhibits nuclear entry by an unknown mechanism.

We recently demonstrated that HIV-1 cores are elastic and that HIV-1 mutants with inelastic capsids are impaired for nuclear entry and infection of nondividing cells. Here we show that CypA prevents infection of nondividing cells by such mutants and inhibits their entry into the nucleus. CypA binding to mutant cores further reduced their elasticity *in vitro*, and this effect was reversed by suppressor mutations that restored nuclear entry. We suggest that HIV-1 nuclear entry involves temporal modulation of capsid elasticity by host proteins prior to and during traversal of the nuclear pore.

---

An obligate step in infection by retroviruses is integration, which requires access to host cell chromatin residing within the nucleus. As a lentivirus, HIV-1 can enter the nuclei of nondividing cells, including resting T cells and terminally differentiated macrophages. This ability is endowed by the viral capsid (1, 2), a closed lattice of hexamers and pentamers forming a shell encasing the viral genome and enzymes required for synthesis and integration of proviral DNA. During early stages of infection, the capsid promotes viral reverse transcription by sequestering the internal components of the viral core including reverse transcriptase, the nucleocapsid protein, and the viral RNA genome (3–6). The capsid also interacts with components of the nuclear pore complex and passes through it while remaining mostly or fully intact (7–9). While the detailed mechanism by which traversal of the nuclear pore complex by the megadalton-sized viral core occurs is unknown, recent evidence indicates that the capsid serves as a nuclear transport receptor by forming polyvalent interactions with FG repeat domains of several nucleoporins which form a gel-like plug in the central channel of the nuclear pore (10, 11). Thus, by mimicking nuclear import receptors, the capsid may penetrate the plug without employing host cell transport factors required for cellular cargo.

In addition to nucleoporin binding, nuclear entry of the HIV-1 core depends on a specific physical property of the viral capsid. Using atomic force microscopy (AFM) to examine the consequences of strong compression of native HIV-1 cores, we recently demonstrated that the HIV-1 capsid is highly elastic and that this property is tightly linked to efficient nuclear entry (12). Amino acid substitutions in CA that inhibited nuclear entry resulted in cores with reduced elasticity (i.e., increased brittleness). HIV-1 pseudorevertants that emerged during cell culture adaptation of two such mutants restored elasticity, linking elasticity to HIV-1 infection of nondividing cells. We also observed that the capsid-targeting inhibitors PF74 and Lenacapavir reduce HIV-1 capsid elasticity, further linking nuclear entry to elasticity.

Interactions of the HIV-1 capsid with host proteins other than nucleoporins may also affect nuclear penetration of the viral core. One such protein, cyclophilin A (CypA), binds to a flexible loop on the surface of the capsid protein (CA). The effects of CypA on HIV-1 infection are numerous and varied (reviewed in (13)). Treatment of target cells with the inhibitor cyclosporin A (CsA) prevents CypA from binding to the incoming viral capsid, preventing infection by inhibiting reverse transcription, nuclear entry, and possibly integration, depending on the cell type used in the experiment. In early studies, selection for resistance to CsA resulted in the identification of two CA substitutions (A92E and G94D) that independently conferred resistance of HIV-1 to CsA (14, 15). Subsequently, an additional substitution (T54A) in a distinct region of CA was shown to exhibit a similar phenotype (16). These mutations do not prevent CypA binding to the capsid, which can be blocked by other substitutions (G89V, P90A) in CA. Further adaptation of CsA-dependent mutants resulted in acquisition of a suppressor mutation encoding the A105T substitution, which restored the ability of the A92E, G94D, and T54A mutants to infect cells in the absence of CsA (17). Thus, the resulting mutant viruses are neither inhibited by CsA nor do they require it for infection.

The mechanism by which binding of CypA to the HIV-1 capsid inhibits infection by some CA mutants is unknown. CypA binding can stabilize the HIV-1 capsid and increase its stiffness, which is a physical property determined by the force required to reversibly deform the surface of an object under slight compression (18, 19). However, neither of these effects has been linked to CypA inhibition of infection. In this study, we tested the hypothesis that CypA binding to the viral capsid reduces its elasticity, thereby inhibiting nuclear entry. We observed that CypA reduced the elasticity of wild type HIV-1 cores in vitro in a concentration-dependent manner. HIV-1 mutants with intrinsically inelastic capsids were rendered even more brittle by addition of CypA. Ablation of CypA binding to the viral capsid in target cells restored the nuclear entry of HIV-1 mutants with inelastic capsids. Moreover, the A92E mutant, which is inhibited for infection of nondividing cells by CypA, was rendered highly brittle upon addition of CypA in vitro, and this effect was abolished by addition of the A105T suppressor. We also found that A105T enhanced the infectivity of the E45A mutant, particularly in nondividing cells, and addition of the E45A suppressor mutation R132T to A92E and T54A mutants rendered them able to infect nondividing cells. Collectively, our results support the hypothesis that CypA inhibits nuclear entry of some HIV-1 mutants by further reducing the elasticity of their capsids.

## Results

### Infection of nondividing cells by HIV-1 mutants with inelastic capsids is restored by inhibition of CypA

Yamashita and coworkers previously reported that the HIV-1 mutants E45A and Q63A/Q67A, which are impaired for infection of nondividing cells, can be rescued by addition of CsA (2). These observations suggested that nuclear entry of these mutants is inhibited by binding of CypA to the mutant capsids. We confirmed the earlier results and extended the analysis to two additional mutants with impaired nuclear entry: E180A and E212A/E213A (12). We previously showed that viral cores purified from each of these mutants exhibited an elevated breakage upon strong physical compression in vitro, indicative of reduced elasticity. As previously observed, infection of Hela cells by each of these mutants was significantly reduced by pre-treatment of the cells with aphidicolin, which causes cell cycle arrest at the G1/S phase (Fig. 1A). We also observed that addition of CsA significantly increased infection by each of the mutants (Fig. 1B), particularly in arrested cells.

**Fig. 1.**
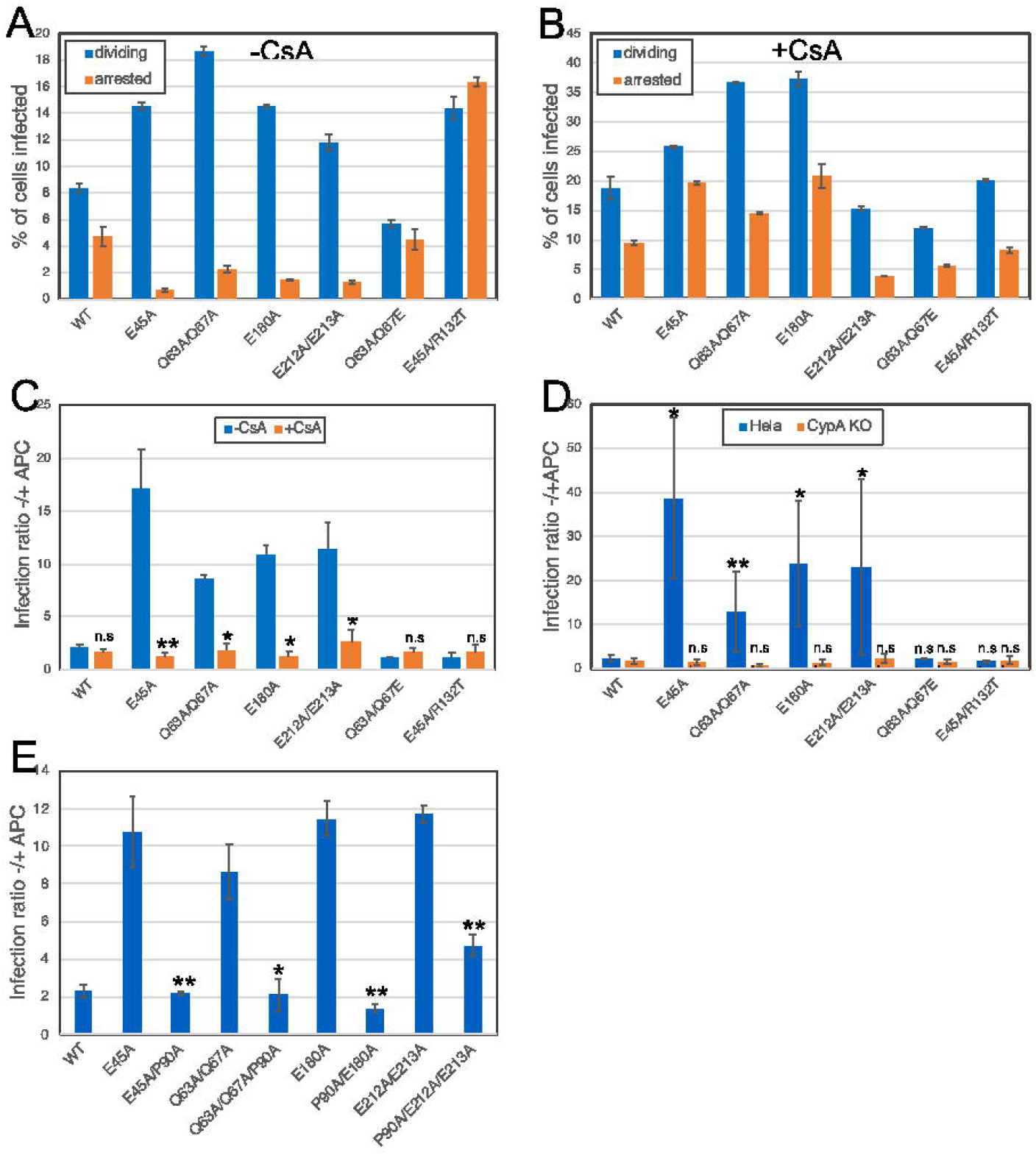
Binding of CypA lo the viral capsid inhibits HIV-1 infection of nondividing cells by HIV-1 mutants with inelastic cores. Infection of wild type (A and B) and CypA KO Hela cells (D) with HIV-1 reporter viruses expressing GFP was quantified by flow cytometry. Cells were seeded in the presence and absence of aphidicolin to induce cell cycle arrest. In panel B, cells were infected in the presence of CsA. Graphs in A and B show results from one representative of three experiments, with average values from duplicate infections. Error bars show the range of the two values. Panel C shows the infection ratios on dividing vs. nondividing cells in the presence and absence of CsA. Mean values from four independent experiments are shown, with error bars representing standard deviations. Symbols represent results of statistical analysis of the effect of CsA for each virus. In panel D, the effects of CypA expression on infection of nondividing cells are shown. Statistical symbols represent comparisons between values from mutant and wild type viruses. Results shown in panel E are from viruses encoding the P90A substitution to prevent binding of CypA to the viral core in target cells. Values shown are means from three independent experiments, with error bars representing standard deviations. Significance was evaluated comparing each mutant with and without the P90A substitution using a paired ratio T test in Graphpad Prism.*: p<0.05; **: p<0.005.

To confirm that CypA binding to the viral capsid preferentially inhibits infection of arrested cells by the mutants, we analyzed the infection of dividing and arrested Hela cells ablated for CypA expression. Comparing the ratio of infection of dividing vs. arrested cells for each virus, we observed a significant reduction in infection of nondividing cells for each of the four mutants (E45A, Q63A/Q67A, E180A, and E212A/E213A) vs. the wild type (Fig. 1C). By contrast, in CypA knockout cells, a marked rescue of infection of nondividing cells was observed for each mutant virus, resulting in no significant difference in cell-cycle dependence of infection vs. the wild type (Fig. 1C). Importantly, infection by the two pseudorevertants E45A/R132T and Q63A/Q67E behaved like the wild type, in that no sensitivity to inhibition by cell cycle arrest was observed in both wild type and CypA knockout cells (Fig. 1, A and C). We previously showed that these two mutants exhibit improved elasticity relative to their parental mutants, suggesting that reduced capsid elasticity renders HIV-1 mutants susceptible to inhibition by target cell CypA.

To confirm that CypA binding to the viral capsid is responsible for inhibition of infection of nondividing cells, we constructed mutants bearing both the elasticity reducing mutations and the P90A substitution, which prevents CypA from binding to the capsid. The addition of the P90A substitution significantly increased the ability of the inelastic capsid mutants to infect nondividing cells, resulting in a reduced infection ratio in untreated and aphidicolin-treated cells (Fig. 1D). Collectively, these results establish that CypA binding to the viral capsid selectively inhibits infection of nondividing cells by these HIV-1 mutants containing inelastic capsids.

### CypA inhibits nuclear entry of HIV-1 mutants with inelastic capsids

To test whether CypA inhibits infection of nondividing cells by preventing HIV-1 nuclear entry, we observed reduced accumulation of 2-LTR circles and late reverse transcription products in arrested vs. dividing Hela cells by wild type HIV-1 and the four mutants, with the mutants exhibited greater reductions than WT in arrested cells. Parallel assays in CypA-deficient cells revealed a marked increase in nuclear entry by the mutants as reflected in increased ratios of 2-LTR circles to late reverse transcripts (Fig. 2A). As a complementary approach, we also examined nuclear entry by tracking the intracellular location of integrase-neonGreen (INmNG)-labeled HIV-1 cores in cells by confocal microscopy. Nuclear entry of the E45A, E180A, and E212A/E213A mutants was markedly increased in cells lacking CypA (Fig. 2B). Surprisingly, while the results of the 2-LTR circle assays indicated increased nuclear entry of Q63A/Q67A in CypA-deficient cells (Fig. 2A), the imaging approach failed to detect nuclear entry by this mutant under any of the conditions (data not shown). We have not established the cause of this apparent discrepancy, but it is possible that the Q63A/Q67A cores exhibit unstable association of the labeled IN protein during reverse transcription. Nonetheless, these results show that CypA binding to the HIV-1 capsid inhibits infection of nondividing cells by preventing nuclear entry of the viral core.

**Fig. 2.**
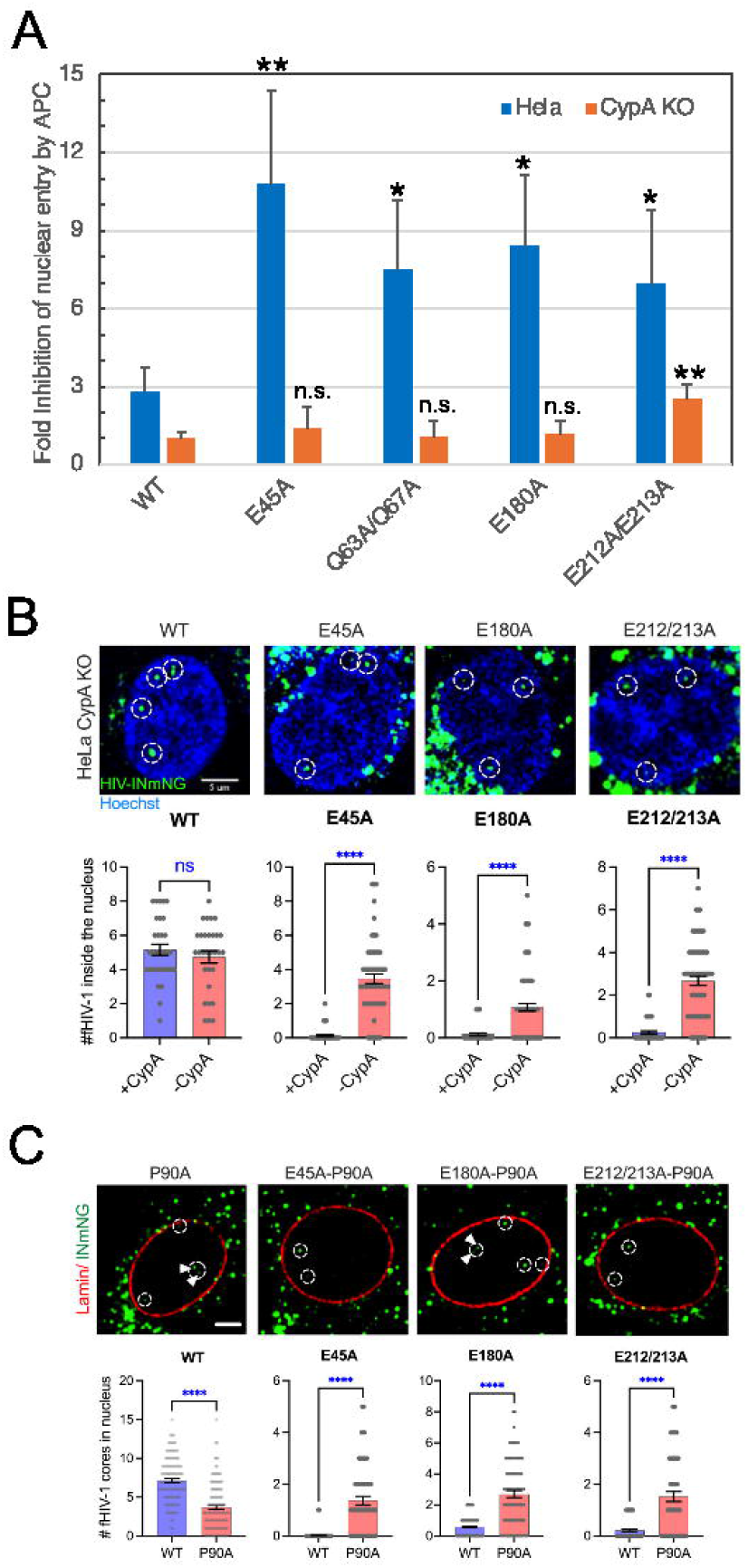
CypA inhibits the nuclear entry of HIV-1 mutants with inelastic capsids. Nuclear entry was monitored in aphidicolin-treated Hela cells by **(A)** quantifying the ratio of 2-LTR circles to late reverse transcripts (second strand transfer DNA) by qPCR, or by **(B)** confocal imaging of INmNG-labeled fluorescently HIV-1 core (fHIV-1) in the nucleus at 8 hours post infection. **(C)** The nuclear entry of inelastic capsids bearing the P90A substitution was determined in emiRFP-670 tagged lamin-81 expressing TZM-bl cells. Images of fHIV-1 nuclear entry in aphidicolin-arrested Hela CypA-KO or P90A-mutant infections in WT-TZM-bl cells are shown. Results were compiled from four independent experiments, with mean values and standard deviations (A), or standard errors of the mean (B and C) shown. Statistical significance was determined using Mann-Whitney Rank Sum test, p-values >0.5 not significance (ns), and <0.0001 (highly significant) is marked by^****^

### Binding of CypA to the viral capsid reduces the elasticity of native HIV-1 cores

In principle, coating of the viral capsid by CypA could inhibit nuclear entry sterically by increasing the effective size of the core. However, the ability of suppressor mutations distal to the CypA binding site to rescue nuclear entry in nondividing cells suggested the possibility of a different mechanism. We previously showed that the suppressors reversed the reduced elasticity of the original mutants, suggesting that CypA binding may further alter the elasticity of the capsid. To test this, we employed atomic force microscopy imaging to examine the state of HIV-1 cores following a strong forced compression (reflecting the elasticity of the capsid). To quantify elasticity, individual viral cores were subjected to strong forced compression, and the fraction of cores that undergo permanent structural damage was determined by AFM imaging. Addition of CypA increased the breakage of wild type cores in a concentration-dependent manner, resulting in 30% of cores undergoing breakage at a CypA concentration of 40 μM (Fig. 3A). As a control for potential nonspecific effects of the recombinant CypA protein, we also tested its effect on HIV-1 cores bearing the P90A substitution in CA, which abolishes CypA binding. Intriguingly, the mutant exhibited approximately 1.5-fold increased breakage vs. the wild type in the absence of CypA (Fig. 3A). Addition of CypA had no effect on the breakage frequency (Fig. 3A), thus confirming that binding of CypA is required for its effects on the elasticity of HIV-1 cores.

**Fig. 3.**
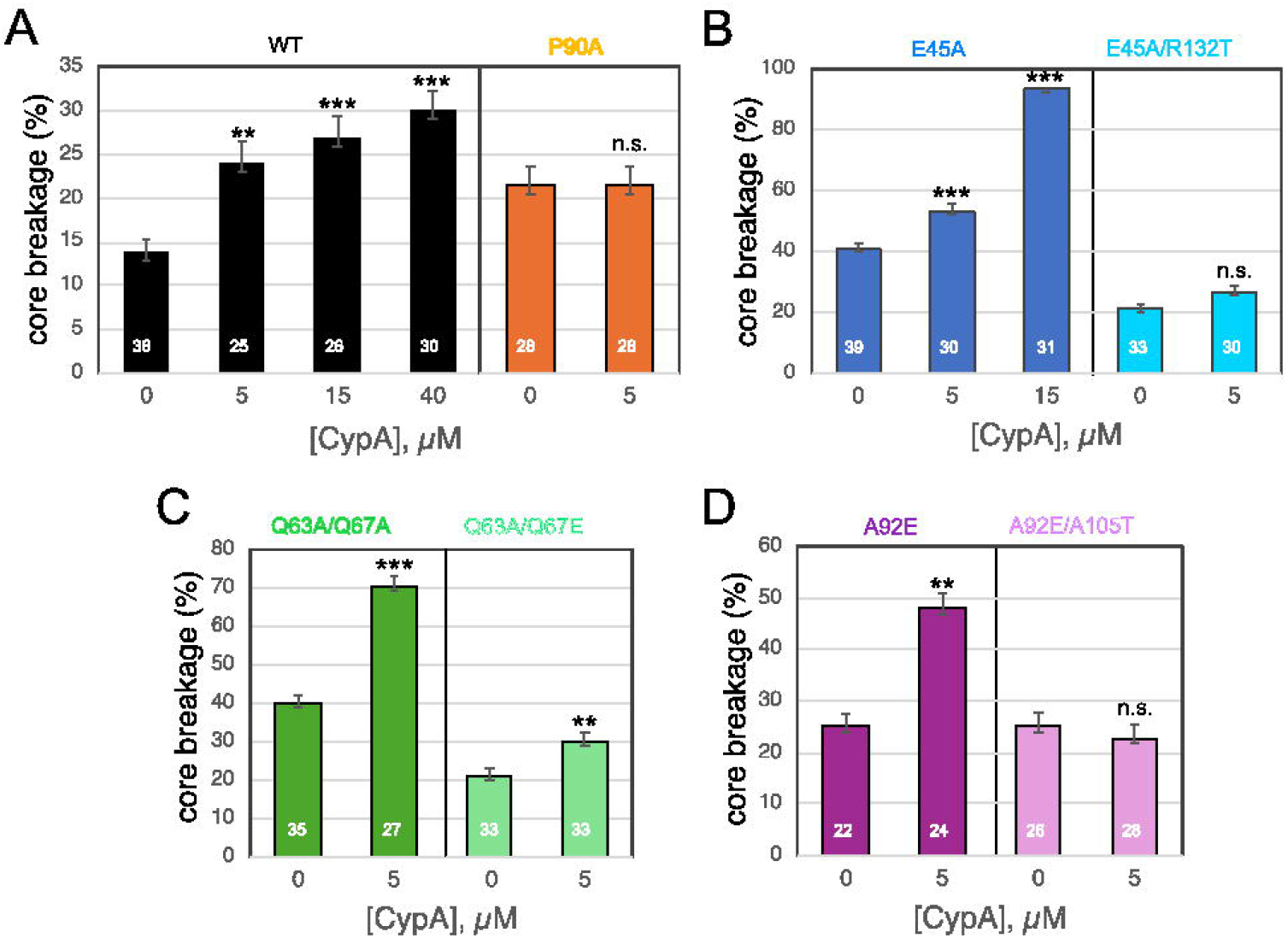
Effects of CypA on the elasticity of purified HIV-1 cores. Viral cores were purified from detergent-treated virions in the presence of IP6 and immobilized on glass slides under native (unfixed and unstained) conditions. Cores were first identified by AFM scanning. A series of cores was then individually subjected to imaging by scanning, compressed by application of the probe at high force, allowed to recover, and rescanned. Broken cores were identified as visible damaged (decreased height and/or altered shape). The analysis was subsequently performed on a separate set of attached cores treated with the indicated concentrations of CypA. Results shown are the percentage of cores that underwent breakage upon CypA addition, with error bars depicting the standard error of the mean. The number of cores analyzed for each condition is shown within the corresponding bar. Panel (A): wild type and P90Acores; (B): E45Aand E45A/R132T cores; (C): Q63A/Q67A and Q63A/Q67E cores; (D): A92E and A92E/A1OST cores. The error of the mean in was obtain by running a bootstrap analysis. Significance of the effect of CypA (relative to the respective no-CypA value) was evaluated using a Chi-squared test for independence. n.s.: p>0.05; **: p<0.01; ***: p<0.001. Data from measurements of WT, E45A, E45A/R132T, Q63A/Q67A and Q63A/Q67E without CypA were taken from Deshpande et al. (12).

CypA increased the breakage of E45A cores from an initial level of 41% (no CypA) to 53% (5 μM CypA) and 93% (15 μM CypA) (Fig. 3B). Upon addition of 15 μM CypA, a large fraction of the cores underwent breakage even prior to AFM compression. Following treatment of immobilized E45A cores with 40 μM CypA, no intact cores were observed (data not shown). Thus, at high CypA concentrations E45A cores appeared to undergo destruction, consistent with a previous study reporting disruption of assembled CA tube structures by 40 μM CypA (19). Collectively, these results indicate that CypA decreases the elasticity of the E45A mutant HIV-1 capsid, resulting in capsid breakage at high concentrations.

As previously reported (12), cores purified from the E45A pseudorevertant E45A/R132T exhibited a breakage frequency approximately half that of E45A mutant cores (Fig. 3B). Addition of CypA to E45A/R132T cores increased their brittleness only slightly, resulting in a core breakage value similar to that of CypA-bound wild type cores (Fig. 3B).

As with E45A, addition of CypA further increased the breakage of intrinsically inelastic Q63A/Q67A mutant cores (Fig. 3C). By contrast, cores from the Q63A/Q67E mutant exhibited elasticity values approximately half that of the Q63A/Q67A and were less affected by CypA (Fig. 3C). Together, these results indicate that CypA inhibition of nuclear entry is linked to its effects on HIV-1 core elasticity.

Given the similarity between the effects of CypA on infection by inelastic capsid mutants and its effects on CsA-dependent mutants, we also tested the elasticity of purified A92E mutant cores. In the absence of CypA, approximately 25% of A92E cores underwent breakage upon compression (Fig. 3D), suggestive of elasticity between that of wild type and E45A mutant cores. Addition of 5 μM CypA increased the breakage frequency to 48%—roughly double that of wild type cores. By contrast, the elasticity of cores purified from the A92E/A105T pseudorevertant was unaffected by CypA (Fig. 3D). These results suggest that the impaired elasticity of the A92E mutant capsid and the further reduction by CypA are related to the impaired ability of the A92E mutant virus to infect nondividing cells.

### The E45A substitution does not detectably alter the affinity of CypA for the viral capsid, while the A92E substitution leads to 2-fold increase in the dissociation constant

The E45 side chain is located at an intersubunit interface in the capsid hexamer, while A92 lies within a flexible loop that is exposed on the outer surface of the capsid lattice and to which CypA binds (model shown in Fig. 4A). CypA binding to the HIV-1 capsid has been reported to result in helical array of protein on the capsid surface, and binding could be cooperative and affected by changes in lattice properties (20, 21). To determine whether changes in capsid-binding binding affinity may account for the inhibition of nuclear entry by CypA, we tested whether E45A and A92E substitutions affect CypA binding to HIV-1 cores. Virions were immobilized on glass coverslips and permeabilized with the pore-forming protein Streptolysin O followed by addition of fluorescently labeled CypA protein (Fig. 4B). Binding of CypA to cores was quantified via TIRF microscopy (22). Fitting of the binding data measured at a range of CypA concentrations with an equilibrium binding model yielded a dissociation constant (K_d_) of 25.2 μM for cores with wild type CA (Fig. 4C). As expected, the K_d_ measured with CA E45A cores (27.9 μM) was comparable to the value for the wild type (Fig. 4D). By contrast, the A92E substitution led to a modest decrease in CypA affinity to the core (56.5 μM; Fig. 4E). This 2-fold increase in K_d_ was consistent with the effect of other substitutions at A92 on CypA binding measured by surface plasmon resonance (wild type CA, 15+/−5 μM; CA A92G, 32 μM; CA A92V, 22 μM) (23). At the estimated CypA concentrations in the cell (10-30 μM), this decrease in affinity would lead to a reduction in CypA binding to the core of 35-50%. Thus, the effect of the A92E substitution on CypA affinity for the viral capsid does not appear to account for the observed inhibition of infection by the host protein.

**Fig. 4.**
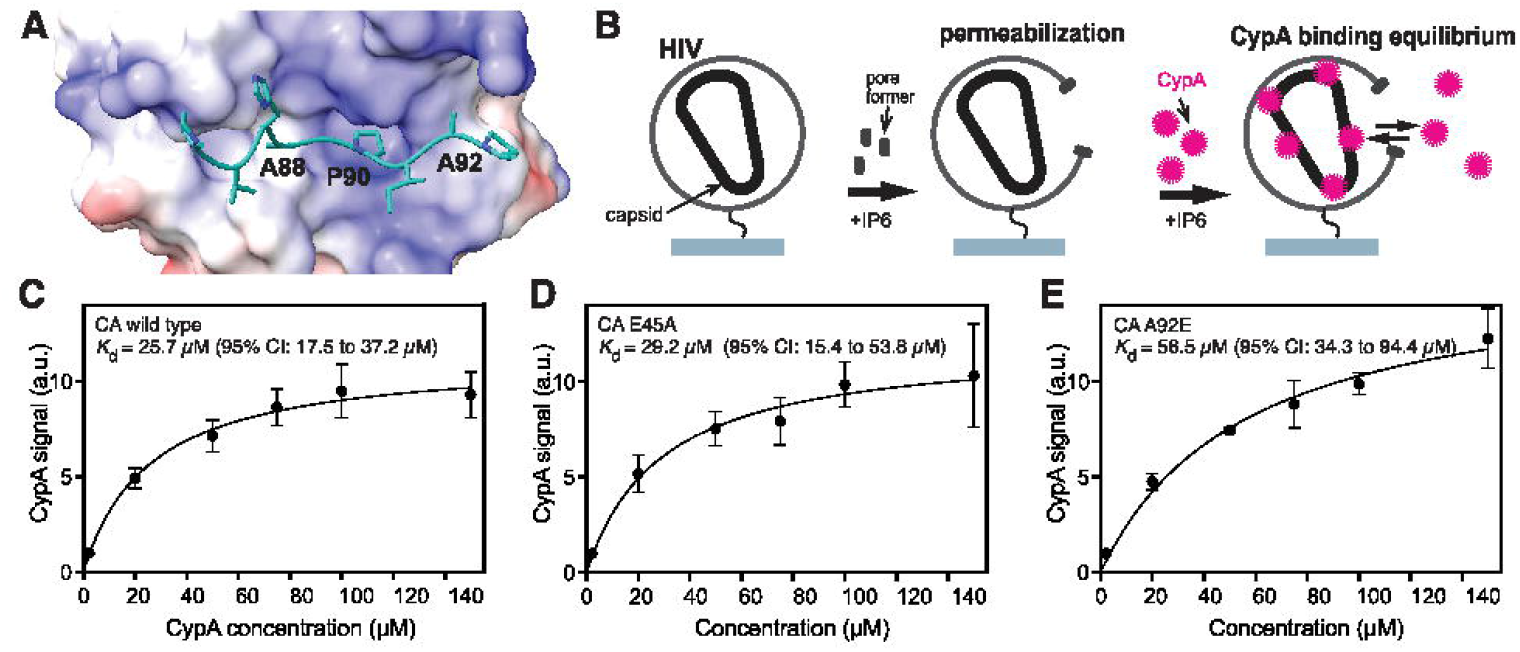
CypA affinity for HIV-1 cores in vitro is not affected by the E45A substitution, but the A92E substitution leads to a 2-fold increase in the dissociation constant. (A): Structural model for the CypA binding loop of HIV-1 CA bound to CypA. (B): Diagram illustrating the experimental approach. Virions were immobilized onto glass, permeabilized with Streptolysin 0, and incubated with various concentrations of fluorescently labeled CypA. Binding of CypA to the viral capsid is detected and by TIRF microscopy and quantified by image analysis. (C-E): Median CypA signal (filled circles) bound at equilibrium to cores in permeabilized HIV with wild type CA (C), CA E45A (D), and CAA92E (E) measured by TIRF microscopy as a function of CypA concentration and fit of an equilibrium binding model (black line).

### The CA suppressor mutations A105T and R132T rescue infection of nondividing cells by the E45A mutant and by CsA-dependent mutants

Our results reveal a phenotypic similarity between the inelastic capsid mutants E45A, Q63A/Q67A, E180A, and E212A/E213A and the CsA-dependent mutants T54A, A92E, and G94D. Because the E45A and the A92E infection phenotypes and CypA-dependent elasticity impairments were rescued by distinct suppressor mutations, we asked whether the suppressors could act reciprocally to rescue infection of arrested Hela cells. To this end, we created mutants bearing the reciprocal suppressors (E45A/A105T; T54A/R132T, and A92E/R132T) and assayed them for infection of arrested cells. We observed enhanced infectivity of each double mutant relative to its parent (Fig. 5A). In arrested cells, the mutants also exhibited improved infection relative to their parental viruses, such that the overall inhibition by aphidicolin treatment was significantly reduced by the suppressors (Fig. 5B). These results demonstrate that the CsA-dependent mutant suppressor A105T increases the infectivity of the inelastic capsid mutant E45A and reverses its inability to infect nondividing cells. Reciprocally, the E45A suppressor mutation R132T rescued infection of nondividing cells by the CsA-dependent mutants T54A and A92E. These results demonstrate that suppressor mutations that reverse capsid elasticity defects also restore the ability of HIV-1 mutants to infect nondividing cells.

**Fig. 5.**
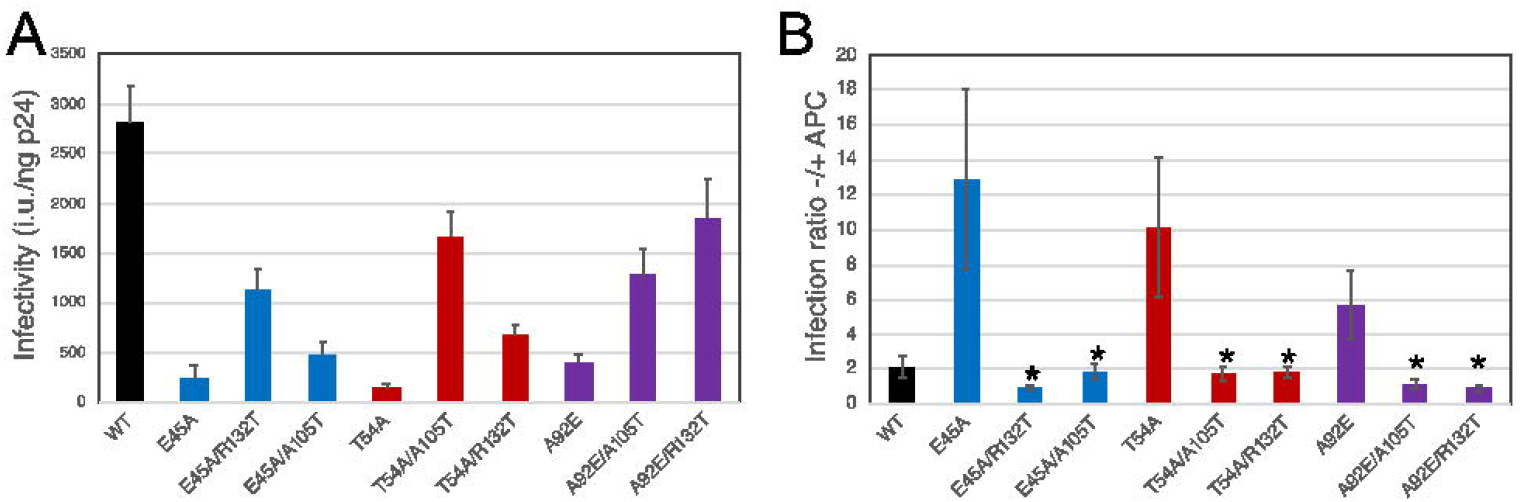
Reciprocal rescue of nondividing cell infection by suppressors of inelastic and cyclosporin A-dependent mutants. Infection of dividing and arrested Hela cells by the indicated HIV-1 mutants was quantified by flow cylometry. (A): lnfectivity of the viruses on dividing cells; (B): ratio of infection on dividing and arrested cells. Asterisks shown in panel B indicate significant differences (p<0.05) between the infection ratio for each double point mutant relative to its respective single mutant. Shown are the mean values of results from three independent experiments with error bars representing one standard deviation.

## Discussion

In this study, we show that CypA, a protein that binds to the HIV-1 capsid in the cytoplasm, can inhibit infection of nondividing cells by preventing nuclear entry. Through the analysis of HIV-1 mutants bearing substitutions in the capsid protein, we linked CypA inhibition of nuclear entry to its effect on the elasticity of the viral capsid. For mutants with intrinsically brittle capsids (E45A and Q63A/Q67A), CypA reduced their elasticity in an additive manner. Suppressor mutations reversed the effects of CypA on elasticity and restored infection of nondividing cells. Thus, binding of CypA to the viral capsid inhibits the ability of HIV-1 mutants to infect nondividing cells by reducing capsid elasticity.

For the four mutants previously reported to contain inelastic capsids, CypA inhibited nuclear entry. By contrast, cores from A92E mutant virus exhibited an intermediate elasticity that was further reduced by CypA. A92E and several other CsA-dependent mutants are apparently competent for nuclear entry despite their impaired ability to infect nondividing cells (17, 24). Thus, the infection barrier imposed by CypA on A92E mutant appears to be at a step following nuclear entry—possibly integration. CypA has previously been shown to alter HIV-1 integration targeting (25, 26), consistent with the host protein affecting viral capsid function following nuclear entry. Collectively, our results suggest that a threshold level of capsid elasticity is required for efficient HIV-1 nuclear entry.

Recent groundbreaking studies have observed intact or nearly intact HIV-1 cores within the cell nucleus, challenging an earlier view that disassembly of the viral capsid must occur in the cytoplasm to permit nuclear entry (9, 27, 28). Moreover, CypA bound to the viral capsid in the cytoplasm apparently dissociated from the core prior to or during passage through the nuclear pore (28). Our results are compatible with this view. While we observed that the inhibition of nuclear entry by CypA was most pronounced for mutants with intrinsically inelastic capsids, we also detected a small extent of inhibition of nuclear entry of wild type HIV-1 and a reduction in wild type core elasticity. These observations, coupled with our finding that CypA binding reduced the elasticity of mutant HIV-1 cores, suggest that CypA regulates either the timing or efficiency of nuclear entry by modulating the elasticity of the capsid. Accordingly, CypA was previously observed to slow the nuclear entry of wild type HIV-1 using cell imaging (29).

The inhibitory effects of CypA on HIV-1 replication can vary substantially with the cell type used for infection. Our work employed Hela cells that are frequently used in studies of HIV-1 infection. Future work will be required to determine whether the same effects are observed in relevant primary cell types including CD4+ T cells and macrophages. The A92E mutant studied here was originally identified by selection for CsA resistance in primary human T cells, and was found to be dependent on the drug, suggesting that CypA is inhibitory to the mutant in relevant cells.

We also observed that the R132T suppressor rescued infection of arrested cells by CsA-dependent mutants A92E and T54A; reciprocally, the A105T suppressor of CsA-dependent mutants rescued E45A infection of arrested cells. These suppressors restored elasticity to their parent mutants, suggesting that HIV-1 adapts to drug selection and host cell pressure by passing through fitness troughs involving changes in capsid properties, including elasticity. We previously showed that the capsid-binding inhibitors PF74 and Lenacapavir reduce HIV-1 capsid elasticity at concentrations of the compounds that inhibit nuclear entry of the viral core (12). Inhibition of HIV-1 infection by PF74 is enhanced by binding of CypA to the viral capsid (30), which may be linked to CypA’s effect on elasticity. We suggest that HIV-1 may develop resistance to capsid inhibitors by altering capsid elasticity.

Our study further demonstrates the importance of capsid elasticity in HIV-1 nuclear entry and establishes that it is controlled by the binding of a host protein. However, our work does not identify the mechanism by which core elasticity promotes traversal of the nuclear pore complex. One possibility involves compression being exerted on the core during penetration of the nuclear pore owing to the presence of the high density of nucleoporin side chains in the central plug. Supporting this hypothesis is the observation that the nuclear pore complex can undergo structural damage (i.e., “cracking”) during HIV-1 nuclear entry, suggesting that strain may be experienced by the core during penetration of the pore (27, 28). The dynamic interactions of the capsid with FG repeat polypeptides occurring during nuclear pore penetration may further alter the properties of the capsid in a manner that is not yet understood. Finally, capsid-binding host proteins that inhibit HIV-1 nuclear entry, including Mx2, CPSF6-358, and TRIM5α, may do so by modulating capsid elasticity.

While our results link CypA’s effect on elasticity to its ability to inhibit nuclear entry of HIV-1 mutants, this action may be distinct from the beneficial effect of CypA in wild type HIV-1 infection, which has been attributed to protection of the viral core from restriction by TRIM5α and cloaking of the reverse transcribed viral DNA from cytoplasmic sensing mechanisms (31–33). The apparent shedding of CypA from the capsid during nuclear entry could resolve this dilemma (28). Thus, it appears that the HIV-1 capsid has evolved to engage the host cell protein CypA for protection in the cytoplasm while evading its ability to inhibit nuclear entry. These opposing demands may have contributed to the high sensitivity of the multifunctional HIV-1 capsid protein to mutations (34).

## Online Methods

### Cells and viruses

Hela cells were purchased from the American Type Culture Collection. The clonal cell line Hela-CypA KO were generated by CRISPR knockout of the cyclophilin A gene using the guide sequence cloned into pLenti-CRISPR2, as previously described (35). Depletion of endogenous CypA was demonstrated by immunoblotting of cell lysates. Cells were cultured in DMEM containing 10% fetal bovine serum and penicillin and streptomycin. Aphidicolin and cyclosporin A were purchased from Sigma and dissolved in water and DMSO, respectively. Solutions were stored in aliquots at −80°C and thawed prior to use.

The HIV-GFP reporter viral proviral plasmids encoding substitutions in CA were previously described: E45A, E45A/R132T, Q63A/Q67A, Q63A/Q67E, E180A, E212A/E213A (12). HIV-GFP plasmids encoding E45A/A105T, T54A, T54A/A105T, T54A/R132T, A92E/A105T, and A92E/R132T were constructed by transferring BssHII-SpeI, BssHII-ApaI, and SpeI-ApaI restriction fragments from the corresponding R9 molecular clones, based on the location of the mutations in the HIV-1 genome (16, 17).

Viruses were generated by transfection of 293T cell monolayers in 10 cm dishes using polyethylenimine (36). Supernatants were harvested 36-42h post-transfection, clarified by centrifugation, and passed through 0.45 μm syringe filters. Aliquots were frozen and stored at −80°C until used for infection assays. Infections of Hela cells were performed in 24-well plates, as previously described (12). Extent of infection was determined by flow cytometry for GFP expression in cell suspensions that had been inactivated by chemical fixation. For infections of arrested cells, cells were seeded in culture media containing aphidicolin (2 μg/ml). The virus stocks were diluted in media containing aphidicolin. One day after inoculation, the inoculum was removed by aspiration and the cultures replenished with fresh medium lacking aphidicolin. In experiments involving CsA, the drug was added to the cell culture media to a final concentration of 5 μg/ml.

To measure viral infectivity, viral titers were calculated from the dilution used in the infection and the percentage of infected cells and divided by the p24 concentration of the inoculum. The latter was determined using an in-house ELISA protocol, as previously described (37).

Nuclear entry was assayed by qPCR quantification of 2-LTR circles and late reverse transcripts in purified total cellular DNA, as previously described (12).

### Expression and purification of recombinant HisCypA

N-terminally His-tagged CypA (HisCypA) was recombinantly expressed in E. coli strain C41 (DE3). Cultures were grown in 2×YT medium with ampicillin and induced mid-log phase with 1 mM IPTG for overnight expression at 18°C. Cells were pelleted, resuspended in buffer (50 mM Tris, pH=9.0; 150 mM NaCl; 1 mM DTT; cOmplete™ protease inhibitor; 20% BugBuster (Merck Millipore)), and lysed by sonication. Lysates were clarified, and His-tagged protein was captured by immobilized metal ion affinity chromatography on a 5□mL Ni2+-NTA HP column (Cytiva). Following a wash step (50 mM Tris pH 9.0, 150 mM NaCl, 1 mM DTT, 20 mM imidazole), HisCypA was eluted using buffer containing 300 mM imidazole. To remove nucleic acids, the eluate was passed through a 5 mL Q anion exchange column (Cytiva). Final purification was performed via size exclusion chromatography on a HiLoadTM Superdex 75 pg column (Cytiva) with the final storage buffer (50 mM Tris, pH 9.0; 150 mM NaCl; 1 mM TCEP).

### Quantification of HIV-1 nuclear entry by confocal microscopy

Nuclear entry of HIV-1 capsid mutants was determined as previously described (38, 39). In brief, 8×10^4^ or 5×10^5^ TZM-bl cells that stably express the emiRFP670-laminB1 nuclear envelope marker, or the Hela parental (control) and CypA-knock-out (CypA KO) cells, were plated in a 8-well chambered slide (#C8-1.5H-N, CellVis) for single time-point nuclear import experiments. Aphidicolin (Sigma Aldrich) was added to cells (10 µM final concentration) to block cell division. About 14h later, the cells were infected (MOI 1 for nuclear entry) with wild type and mutant HIVeGFP particles fluorescently tagged with Vpr-integrase-mNeongreen (INmNG) to label the viral core. Virus binding to cells was augmented by spinoculation (1500×g for 30 min, 16°C), and virus entry was synchronously initiated by adding pre-warmed complete DMEM medium containing aphidicolin (10 µM) to samples mounted on a temperature- and CO_2_-controlled microscope stage. 3D confocal imaging was carried out on a Leica SP8 LSCM using a C-Apo 63x/1.4NA oil-immersion objective. Tile-scanning was employed to image multiple (4×4) fields of view. For assessing HIV-1 nuclear entry in a fixed time-point (8 hpi), stringent imaging conditions were used i.e., 4x line-averaging, with 0.12 μm/pixel sizes and 0.5 μm spaced z-stacks. 488 and 633 nm laser lines was used to excite the INmNG and emiRFP670-Lamin B1 fluorescent markers, and their respective emission was collected between 502-560 nm and 645-700 nm using GaSP-HyD detectors. For Hela cell experiments, the nuclei were labeled with Hoechst dye and imaged using 405 nm lasers, and emission was detected using GaSP-HyD detectors set at 420-480 nm. 3D-image series were processed off-line using ICY image analysis software (http://icy.bioimageanalysis.org/) (40). HIV-1 nuclear entry in fixed time-point 3D z-stack images was analyzed using an in-house script in the ICY protocols’ module as described in (38).

### AFM assay of HIV-1 core elasticity

Samples for AFM measurements and analysis were prepared as previously described (41). Briefly, 10 µL of isolated HIV-1 cores were incubated for 45 min at room temperature on hexamethyldisilazane (HMDS)-coated microscope glass slides (Sigma-Aldrich) in a mildly humid chamber. AFM measurements were performed on the adhered sample without fixation. Each experiment was repeated at least three times, each time with independently purified pseudoviruses. IP6 was purchased from Sigma-Aldrich (P8810). PF74 (PF-3450074) and Lenacapavir (GS-6207, Gilead Sciences) were purchased from MedChemExpress (MCE), USA. All measurements were carried out with a JPK Nanowizard Ultra-Speed atomic force microscope (JPK Instruments, Berlin, Germany) mounted on an inverted optical microscope (Axio Observer; Carl Zeiss, Heidelberg, Germany). Silicon nitride probes (mean cantilever spring constant of 0.12 N/m; DNP, Bruker, Germany) were used. Topographic images were acquired using the quantitative imaging (QI) mode at a rate of 0.5 lines/s and a loading force of 300 pN and rendered using the WSxM software (Nanotec Electronica).

Core elasticity was assessed as previously described (12, 42) by initially scanning a small rectangular region on the core body at a low loading force (300 pN). Compression was then applied by rescanning the same region at a maximum loading force of 5 nN, after which the force was reduced back to 300 pN. The entire core was subsequently reimaged at 300 pN to evaluate structural alterations. Structural integrity was determined by comparing pre- and post-compression images, and elasticity was quantified as the percentage of intact versus broken cores following forced compression.

### TIRF microscopy CypA binding assay

HIV particles were produced in HEK-293T cells transfected using PEI with a mixture of the plasmids pCRV1-GagPol and pCSGW. The medium was exchanged 18 hr post transfection and the virus-containing medium was collected 66 hr post transfection and centrifuged (2100 × *g*, 20 min, 4 °C) to remove cellular debris. The supernatant was concentrated using a 100 kDa MWCO centrifugal filter. HIV particles were purified using a Sephacryl S500 size exclusion column and labelled with biotin using NHS ester chemistry (ThermoFisher, EZ-Link™ NHS-LC-LC-Biotin, #21343). The biotinylated HIV were purified using a Sephacryl S500 size exclusion column, concentrated and flash-frozen in liquid nitrogen. TIRF microscopy binding assays were performed using microfluidic channel devices as described previously [PMID: 29877795]. HIV-1 particles were immobilized on PLL-PEG-biotin– modified coverslips, permeabilized in the presence of 0.1 mM IP6 with the pore-forming protein streptolysin O and incubated for 30 min to allow disassembly of improperly assembled cores (which cannot be stabilized by IP6). A solution containing 2 µM AF647-labelled CypA (produced and labelled as described in [PMID: 30947724]) mixed with varying amounts of unlabelled CypA to yield final CypA concentrations of 2–150 µM was then introduced into the flow channel. CypA binding was quantified by measuring the intensity of diffraction-limited AF647-CypA spots colocalizing with HIV-1 particles in TIRF images. A full concentration titration was performed within each flow channel, and each dataset represents at least three independent channels. Image analysis was carried out using software generated in house (https://github.com/lilbutsa/JIM-Immobilized-Microscopy-SuitE).

## Author contributions

CA and IR conceived the study, designed the experiments, analyzed the data, and wrote the paper with help from all the authors. JH performed HIV-1 infection and PCR-based nuclear entry experiments and analyzed the results. AD, YT, and LS prepared HIV-1 core samples and conducted AFM measurements and analysis. NRD assisted with HIV-1 core preparation. ADWD, MLK, and ACF performed imaging-based nuclear entry assays. CMG performed CypA-core binding experiments. JOK and LJ provided recombinant CypA protein. JS generated mutant pCRV1-Gag-Pol plasmids.

